# Negative DNA supercoiling makes protein-mediated looping deterministic and ergodic within the bacterial doubling time

**DOI:** 10.1101/2021.02.25.432970

**Authors:** Yan Yan, Wenxuan Xu, Sandip Kumar, Alexander Zhang, Fenfei Leng, David Dunlap, Laura Finzi

## Abstract

Protein-mediated DNA looping is fundamental to gene regulation and such loops occur stochastically in purified systems. Additional proteins increase the probability of looping, but these probabilities maintain a broad distribution. For example, the probability of lac repressor-mediated looping in individual molecules ranged 0-100%, and individual molecules exhibited representative behavior only in observations lasting an hour or more. Titrating with HU protein progressively compacted the DNA without narrowing the 0-100% distribution. Increased negative supercoiling produced an ensemble of molecules in which all individual molecules more closely resembled the average. Furthermore, in only twelve minutes of observation, well within the doubling time of the bacterium, most molecules exhibited the looping probability of the ensemble. DNA supercoiling, an inherent feature of all genomes, appears to impose time-constrained, emergent behavior on otherwise random molecular activity.

## Introduction

Protein-mediated DNA looping is a ubiquitous regulatory process that compacts the genome and may regulate transcription, make DNA more, or less, accessible for replication, and facilitate repair (1–5). For instance, effectively switching on or off genes of the lac operon in response to the availability of lactose is advantageous to avoid wasted metabolism. One would expect molecules displaying regulatory looping to exhibit behavior typical of the ensemble, and such ergodic behavior should be observable on an appropriate timescale for cellular biochemistry. Yet, in the thermal bath of the eukaryotic nucleus, or prokaryotic nucleoid, molecular motion is random and the behavior of a molecule can only be described in probabilistic terms (6, 7). Single-molecule experimentation allows fluctuation analysis (8–10) and has revealed very heterogeneous looping behavior of molecules (11–20). Measurements have shown that protein-mediated loops form and rupture stochastically, and the lifetimes of such loops span several orders of magnitude. Notably, different DNA molecules in the single molecule measurements behaved very differently during observations over a time span of approximately 30 min, comparable to the doubling time of *Escherichia coli* (*E. coli*) (21), the bacterium in which LacI-mediated DNA looping is physiologically relevant. Such heterogeneity seems ill suited for biochemical reactions presumably designed to achieve a certain regulatory outcome.

Extending the interval of observation might produce a more homogeneous response (22, 23), but other factors might also change looping dynamics to create similar homogeneity within a biologically relevant time interval such as the mitotic cycle. For example, the activity of biological molecules depends on their ionic environment (24) and bacteria like *E. coli* must contend with large variations in salt concentrations (25). In addition, there is evidence for sub-diffusive behavior of biomolecules in cells (26–29). These complicate the identification of factors that tune looping dynamics, but it might be possible to identify such factors as those that produce time-constrained ergodic responses (30, 31).

Nucleoid-associated proteins which are abundant in bacteria and are thought to decorate the bacterial genome, organizing and compacting the DNA (32, 33) are among possible factors. The HU protein, an abundant nucleoid-associated architectural protein seems a likely candidate, since knockouts and mutants profoundly modify the *E. coli* (K12) transcriptome (34, 35) and it has been observed to facilitate protein-mediated looping, although not against tension (36). Alternatively, evidence that supercoiling might be important had already appeared in earlier reports of comparisons of LacI dissociation from linearized or supercoiled plasmids containing the O1 and O2 operators separated by 400 base pairs (37).

In the experiments described below, the distribution of probabilities of DNA looping mediated by the lac repressor was measured versus various concentrations of the HU protein and levels of negative supercoiling. Only supercoiling significantly altered stochastic DNA looping to produce homogeneous behavior among a large set of molecules and on the time scale commensurate with the doubling time of the bacterium.

## Results

Previous work has shown that the probability of LacI-mediated looping in DNA molecules with two LacI binding sites (operators) depends on the protein concentration (36). When LacI concentration is low, neither operator may be occupied by a LacI tetramer, the looping probability is low, so the DNA tether remains extended, and the attached bead exhibits large excursions. When the concentration is too high, both operators become occupied by tetramers which, not being able to bind to each other, cannot form loops. Excursions are large for beads attached to these tethers as well. Only at intermediate concentrations, in which one tetramer may bridge two operators, does the probability of looping increase significantly. When loops form, the tether is less extended, and the excursions of the attached bead are restricted. Average looping probability is indicated by crosses in Figure 1, which summarizes ~30 min-long tethered particle motion (TPM) (17, 38, 39) measurements of LacI-mediated DNA looping between the strong O1 and weaker O2 operators separated by 400 bp (36).

**Figure 1.**
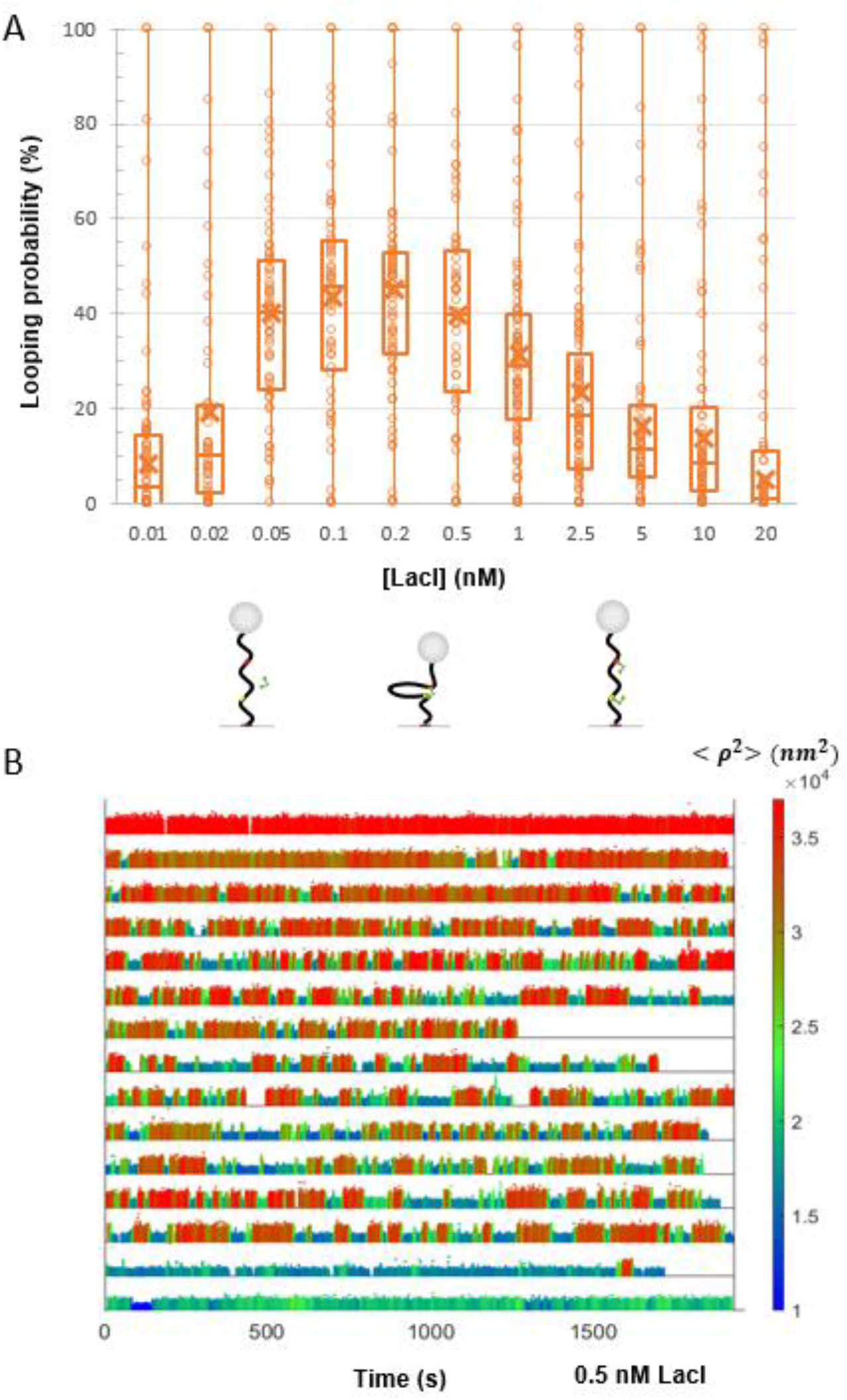
The looping probabilities of different DNA tethers vary widely. (**A**) The calculated looping probabilities of individual tethers exposed to a range of LacI concentrations are summarized in a box-whisker plot. The whiskers span the entire range of probabilities of the ensemble of DNA tethers monitored in each condition. As the LacI concentration was titrated from 0 to 20 nM, the looping probability increased from 0 to approximately 45 and then fell again to 0 when repressor molecules saturated the binding sites. Note however, that at each concentration of LacI, the looping probabilities varied from 0 to 100. The upper and lower borders of the boxes indicate the upper and lower quartiles. The midline and cross of each box indicate the median and average of the distribution of looping probabilities. Schematic diagrams of prevalent DNA/LacI configurations are depicted below their corresponding LacI concentrations. (**B**) Representative temporal records of the TPM excursion parameter < ***ρ***^2^ > with 0.5 nM [LacI] are ranked by looping probability from 0 (top) to 1 (bottom). Unlooped states are depicted in red, and looped states are shown in blue and green corresponding to loops in anti-parallel or parallel configurations respectively. The actual < ***ρ***^2^ > values are encoded using the color scale at right.

Although the average behavior of the population of DNA tethers is clear and follows expectation, the looping probabilities of individual DNA tethers under any given LacI concentration are very heterogeneous. Indeed, the whiskers of each box in the left panel of Figure 1, range from 0 to 100% looping probability. Raw data displaying this behavior is shown in representative temporal records for DNA tethers exposed to 0.5 nM LacI (Figure 1, right panel). Each time record corresponds to a different DNA tether. Unlooped states are depicted in red, and looped states are shown in blue and green corresponding to loops in anti-parallel or parallel configurations (40). Clearly, loop formation and breakdown occur randomly. However, some tethers are never looped, some are always looped, and some toggle between looped and unlooped states with various degrees of probability, calculated as the time spent in the looped state over the total observation time.

To verify that the heterogeneity observed was indeed due to the activity of single lac repressors and not variations among DNA molecules introduced during PCR, a control experiment was performed in which excursions of the tethered beads were monitored for 30 min, and then the first solution containing LacI was washed away with *λ* buffer supplemented with high salt concentration (1 M KCl). To verify washing, the excursions of the same tethered beads were then monitored for 20 min in which no looping was observed. Finally, an identical solution of protein was introduced into the microchamber and excursions of the bead were monitored for another 30 min. For comparison, the percentages of time spent by individual tethers in the looped state during the two observation periods (before and after washout of LacI) were calculated. Supplemental Figure S6 shows the lack of correlation between the looping probabilities measured on individual DNA tethers before and after, suggesting that the looping behavior of a DNA tether is dictated by variations in the activity of associated individual LacI proteins in the two observation periods. While there may be variation in the activity of individual LacI enzymes, the following experiments on supercoiled DNA molecules established conditions in which all DNA tethers exhibited a narrow range of looping probabilities within observations on the timescale of the doubling time of *E. coli*. Thus, activity variations do not appear to be significant.

Such extreme variation in the stability of lac-mediated loops might occur *in vivo*, but it is difficult to rationalize how it could benefit *E. coli* bacteria that should calibrate a response to lactose. Thus, there might be factors *in vivo* that turn looping into a deterministic process. Since protein-mediated looping is a ubiquitous regulatory mechanism across kingdoms, this question transcends the specific organism and is relevant for cells of all organisms. We hypothesized that genome-compacting proteins and DNA supercoiling, which are common to all species, might decrease the variation in looping probability, because they compact DNA and reduce the distance between sites joined by the looping protein. As a model genome-compacting protein, we chose the nucleoid-associated protein HU which is abundant in bacteria, binds non-specifically to DNA, contributes to the overall architecture of the genome, facilitates protein-mediated looping, and influences DNA replication and transcription (36, 41–45).

To determine whether HU can eliminate variation in looping probabilities, tethered particle motion (TPM) experiments were conducted at one LacI concentration (2.5 nM) while the HU concentration was titrated from 0 to ~1 μM. The LacI concentration was chosen such that it would be easy to measure with negligible uncertainty, and increases or decreases in the looping probability would be obvious (36). As shown previously by our lab and others, in the salt condition we use here, the magnitude of excursions of beads tethered to single DNA molecules decreases as HU concentrations increase (36, 46); this protein-induced DNA compaction facilitates looping (36). Indeed, as shown in the left panel of Figure 2, increasing the HU concentration increases the median looping probability driving it from around 20 to 80%. However, the looping probabilities of single DNA tethers ranged from 0 to 100% at each HU concentration as shown by the whiskers in the plot. The representative temporal traces in the right panel of Figure 2, show tethers that never looped, tethers that remained looped throughout the observation, and others that toggled between the looped and unlooped states. This is similar to what was observed for LacI-induced looping without additional factors (Figure 1), indicating that HU, despite its ability to compact DNA and favor looping overall, did not reduce the variation in the looping probabilities of different DNA tethers.

**Figure 2.**
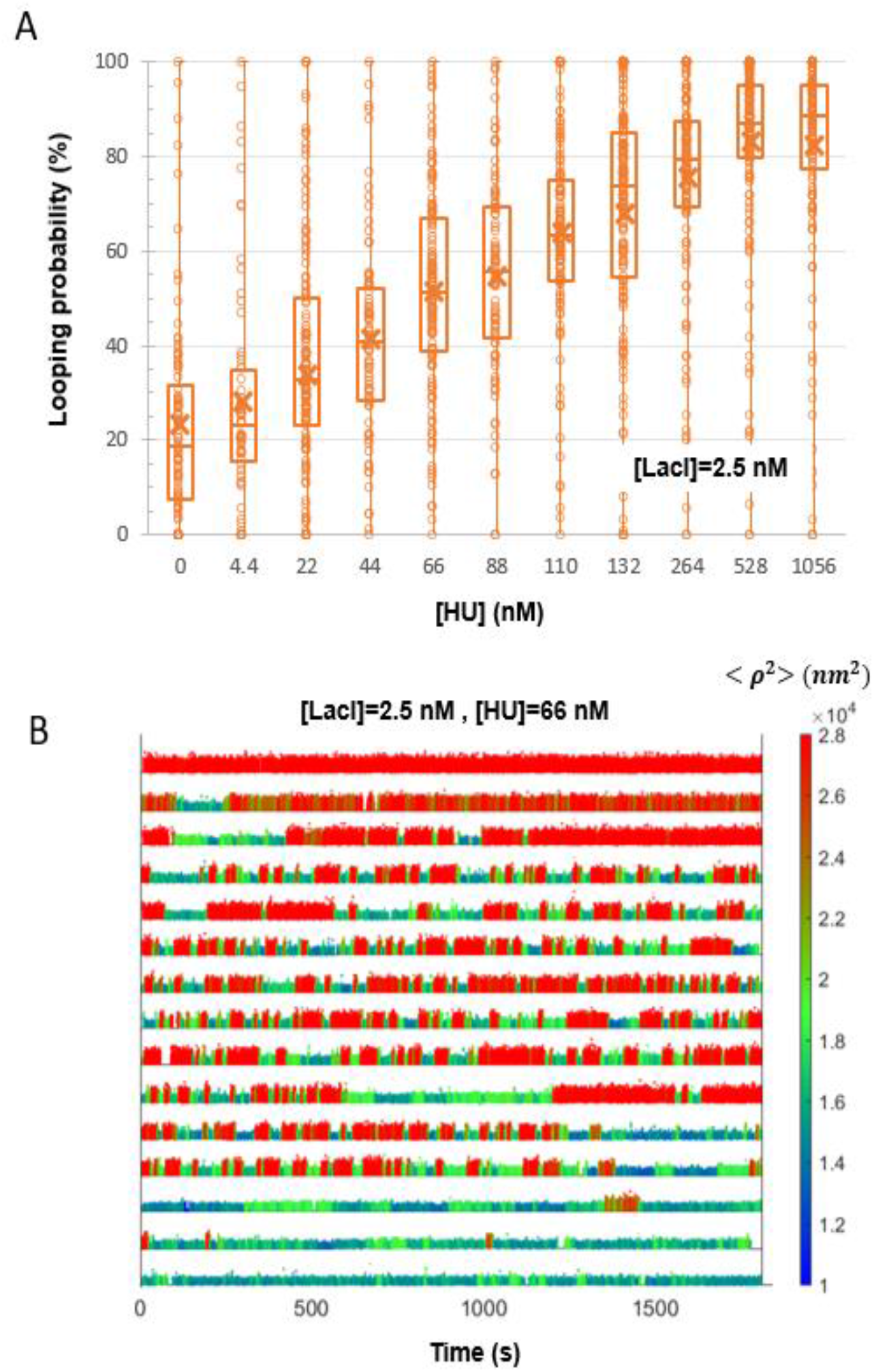
HU did not reduce the variation of looping probabilities amongst different tethers. (**A**) The calculated looping probabilities of individual tethers, exposed to 2.5 nM and a range of HU protein concentrations, are summarized in a box-whisker plot. The whiskers span the entire range of probabilities of the ensemble of DNA tethers monitored in each condition. As the HU concentration was titrated from 0 to 1056 nM, the average looping probability progressively increased from 25 to just above 80. Note however, that at each concentration of HU, the looping probabilities for individual DNA tethers varied from 0 to 100. The upper and lower borders of the boxes indicate the upper and lower quartiles. The midline and cross of each box indicate the median and average of the distribution of looping probabilities. (**B**) Representative temporal records of the TPM excursion parameter < ***ρ***^2^ > with 2.5 nM [LacI] and 66 nM [HU] are ranked by looping probability from 0 (top) to 1 (bottom). Unlooped states are depicted in red, and looped states are shown in blue and green corresponding to loops in anti-parallel or parallel configurations, respectively. The actual < ***ρ***^2^ > values are encoded using the color scale at right.

DNA supercoiling is a second factor responsible for genome compaction in live cells. It refers to the over- or under-winding of a DNA molecule, which is often quantified as the ratio of the number of turns added to a DNA molecule over the number of helical turns in the torsionally relaxed state. In live cells, DNA supercoiling is ubiquitous and dynamic, genomes are, in average, negatively supercoiled, and it has been shown that DNA unwinding under low, physiological forces compacts DNA in a way that facilitates looping by proteins (36). Magnetic tweezers were used to modulate supercoiling and measure its effect on LacI-induced looping, focusing on the level of heterogeneity of the looping probabilities of different tethers. Magnetic tweezers are a convenient tool with which to modulate DNA supercoiling for *in vitro* experiments. Such tweezers utilize a pair of magnets placed close to a microchamber with DNA tethering super-paramagnetic beads to the glass surface. The magnetic field generated by the magnets attracts and orients the beads applying tension to the DNA. In addition, if the magnets are rotated, DNA will be twisted. If DNA is twisted under low tension, either winding or unwinding will induce plectonemes that decrease the extension of the DNA (Supplemental Figure S4).

Figure 3 summarizes measurements of LacI-mediated looping probability for the 2115 and 2011 bp DNA tethers unwound by different amounts under 0.45 pN of tension. Negative supercoiling progressively increased the looping probability (36). The left panel in this figure shows that increased (negative) supercoiling drove the looping probability from 0 to 100%. Note also that the looping probabilities measured for different DNA tethers are tightly grouped around the median values. Negative supercoiling dramatically reduced variation and produced a uniform, deterministic behavior. Similar behaviors were also observed with tensions of 0.25 and 0.75 pN (Supplemental Figure S7). This uniformity is illustrated in the right panel of Figure 3, where individual time traces, recorded under 0.45 pN of tension and −2.5 % supercoiling level, are quite similar and exhibit frequent switching between unlooped (red) and looped (blue/green) states.

**Figure 3.**
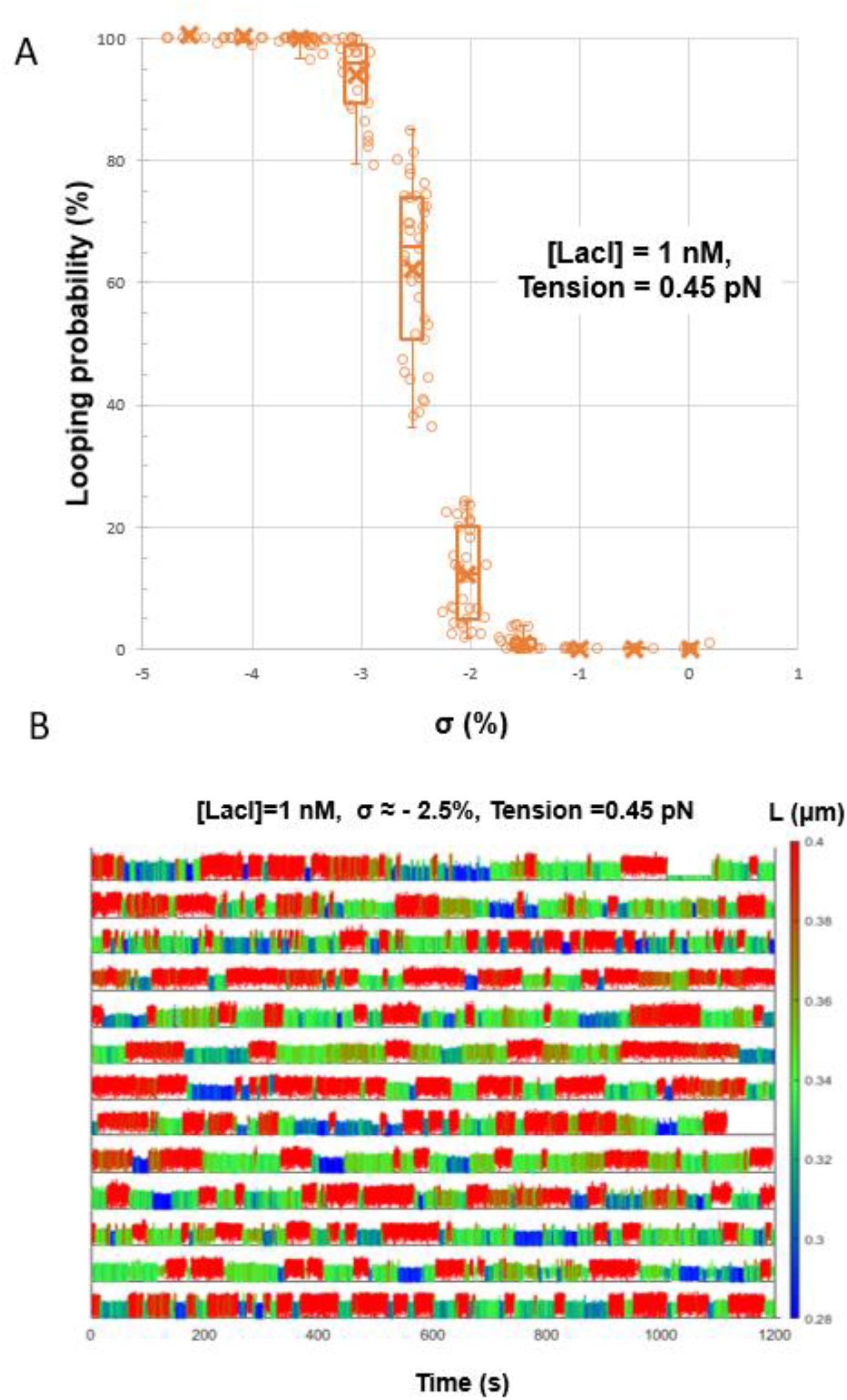
Supercoiling dramatically reduces the variation of looping probabilities amongst DNA tethers. (**A**) The calculated looping probabilities of individual tethers, exposed to 1 nM LacI and negatively supercoiled to varying degrees, are summarized in a box-whisker plot. The whiskers span the entire range of probabilities of the ensemble of DNA tethers monitored in each condition. As the supercoiling was varied from 0 to almost −5% (σ), the average looping probability progressively increased from 0 to 100. Note that at each level of supercoiling, the looping probabilities for individual DNA tethers formed compact distributions. The upper and lower borders of the boxes indicate the upper and lower quartiles. The midline and cross of each box indicate the median and average of the distribution of looping probabilities. The whiskers and quartiles are only distinct for intermediate values of negative supercoiling. For low and high levels of negative supercoiling the whiskers collapse to the median value. (**B**) Representative temporal records of the instantaneous lengths of DNA tethers exposed to 1 nM [LacI] while stretched by 0.45 pN of tension and supercoiled to σ = −2.5% are shown. Unlooped states are depicted in red, and looped states are shown in blue and green corresponding to loops in anti-parallel or parallel configurations respectively. The actual tether length values are encoded using the color scale at right.

## Discussion

### Supercoiling makes protein-mediated looping deterministic

In the absence of supercoiling, the probabilities of LacI-mediated DNA looping for individual DNA tethers with, as well as without, HU range from 0 to 100%. In these conditions, in order for the protein to connect them, the LacI binding sites must juxtapose by 3-D diffusion opposed by a high energy barrier. HU protein helps to overcome this barrier and enhance looping by compacting the DNA to reduce the separation between the operators to be bridged. In contrast, supercoiling induces plectonemes which allow slithering of DNA segments past one another (47, 48). Thus, the operators may juxtapose through 1-D diffusion, across a much lower energy barrier. Negligible bending energy changes accompany slithering and little configurational entropy is lost when LacI connects two sites in a plectoneme. By reducing the dimensionality of the path to juxtaposition, supercoiling produced homogeneous looping probabilities for different DNA tethers that could not be achieved by adding HU to facilitate juxtaposition in three dimensions.

Since HU binding is known to supercoil DNA (49), it might similarly alter looping in a torsionally constrained DNA tether. This was not possible in a TPM experiment (Figure 2) in which single-bond attachments of the DNA to the surfaces would swivel to release any torsion. However, previously published data (36) showed that 1056 mM HU which produces a median value of 85% average looping probability in DNA tethers under no tension (Fig. 2), produced just 30 or 5% in DNA tethers under 0.25 or 0.45 pN of tension, respectively. If HU were actually supercoiling the DNA, then it should have sustained the looping probability under tension. The fact that it did not suggests that HU binding only compacts but does not significantly supercoil DNA in the conditions used here. To verify this, extension versus twist curves were measured for DNA in the presence of HU. In this assay, any unwinding associated with protein or small molecule binding to the DNA will alter the intrinsic twist of the molecule and shift the maximum extension of the molecule with respect to that of bare DNA (50–52). As can be seen in Supplemental Figure S8, increasing the concentration of HU up to 1000 nM steadily contracted a DNA tether stretched by 0.45 pN in 200 mM KCl but negatively supercoiled the molecule by only −1.35 turns. This is equivalent to 0.4% supercoiling, which is insufficient to produce writhe (53). Thus, HU enhances looping by contracting DNA tethers but does not change the dimensionality of looping to promote uniform looping dynamics.

Figure 4 is an illustration of hypothetical energy landscapes for looping, which involves bending and possibly twisting the DNA molecule, as shown in the cartoons on the left. Energy is color-encoded according to the scale at right. The end-to-end distance of the loop segment is represented along the vertical axis. Near-zero end-to-end distance corresponds to the looped state, while the unlooped states are more extended, up to 400 bp. The horizontal axis indicates the supercoiling in the DNA and its handedness. Panel (*A*) describes the experimental conditions in a TPM measurement, with no applied tension or torsion. The DNA may coil, bend, and juxtapose the two operators via 3-D diffusion, as shown in the superimposed cartoon. Different pathways may be followed through the broad and shallow saddle point separating the looped and unlooped states. In panel (*B*), sub-pN tension gently extends DNA in the absence of imposed supercoiling, and the work associated with drawing the two operator sites together increases the energy barrier, effectively attenuating loop formation. These conditions correspond to a magnetic tweezer measurement in which the magnets exert tension but are not rotated to twist the DNA as shown in the superimposed cartoon. However, in plectonemic DNA as shown in the cartoon superimposed on panel (*C*), operator sites can juxtapose as DNA segments slither past each other in the plectoneme. This one-dimensional search for juxtaposition between looped and unlooped states changes the dynamics of looping with respect to the three-dimensional searches required in torsionally unrestrained DNA tethers.

**Figure 4.**
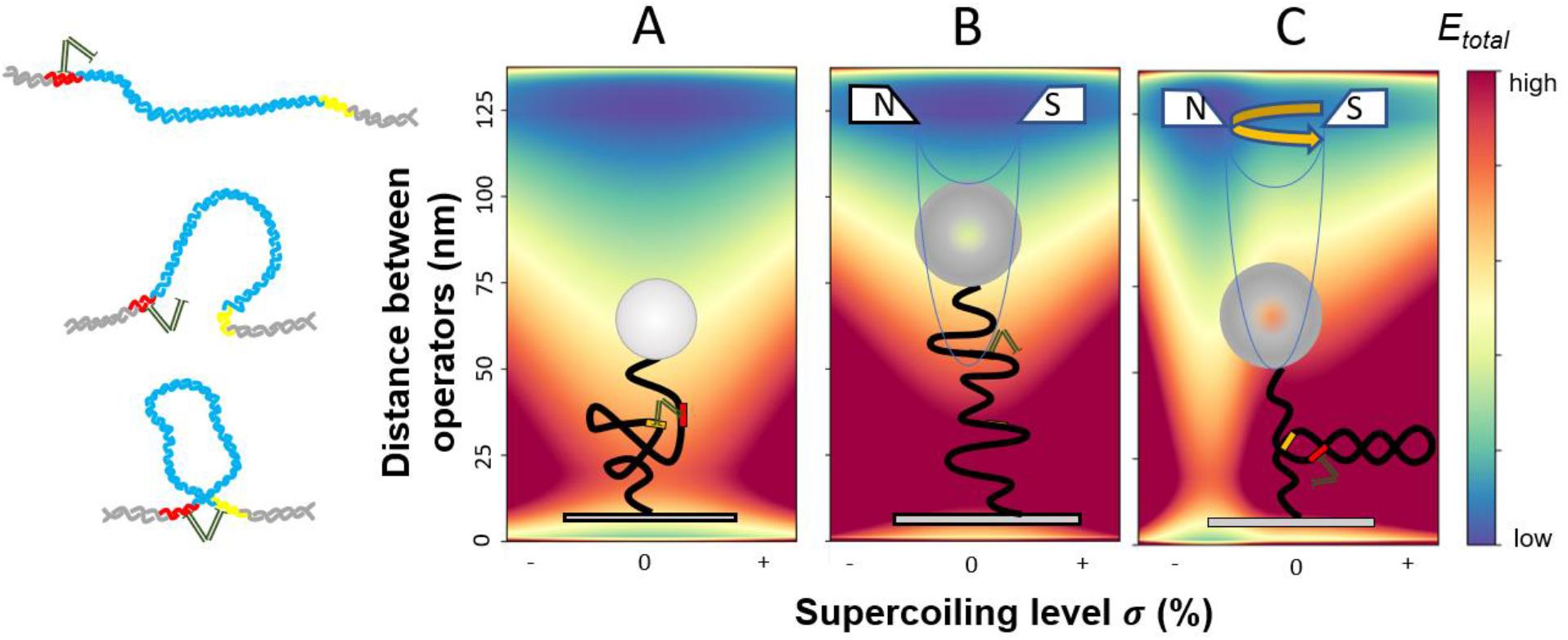
Energy landscapes for loop closure by LacI in different conditions of tension and torsion. The illustrations at the left show possible conformations of DNA tethers corresponding to different separations between the operators. *A-C* are three hypothetical energy landscapes for a DNA tether under different conditions of torsion and tension are represented. The energy values are qualitatively encoded using the color scale at right. The *y*-axis indicates the distance between the protein binding sites that constitute the junction and may vary between zero and 133 nm for a 400 bp DNA segment. DNA supercoiling varies along the *x*-axis. Superimposed on each panel are illustrations of likely DNA conformations under the different conditions of torsion and tension.

### Supercoiling induces ergodicity within a biologically relevant timescale

The TPM records in this work were between 20 and 30 minute-long, slightly below, or equal to the doubling time of *E. coli* which is approximately 30 minutes in the laboratory (21). The heterogeneity of looping probabilities observed in these records without supercoiling were extreme and seemingly at odds with a molecular system designed to respond to the presence of lactose. Such a system was expected to be ergodic, such that sufficiently long observations of single members of the ensemble would have exhibited the statistical behavior of the whole ensemble. Several much longer, five-hour, recordings of LacI-mediated looping in torsionally relaxed DNA were then acquired using TPM (Supplemental Figure S9) and the looping probabilities were measured over temporal windows of different lengths, ranging from 10 min to the entire 5 hour-long recording. Figure 5A is an overlay of the cumulative probability distributions for looping percentages calculated for entire 5-hour records (black) or divided into shorter segments of 10 (blue), 30 (green), 40 (red), 60 (cyan), 80 (magenta), and 100 (yellow) min. Comparison of these distributions shows that recordings greater than or equal to 60 min (cyan) produce distributions like the five-hour (black) distribution in which all average looping probabilities fell between 0-30%. Observations for shorter periods exhibit a tail of high looping probabilities that is not present in distributions from longer observations. Thus, at least 60 min of observation are required to accurately sample the dynamics of LacI-mediated looping in a DNA construct containing the O1 and O2 operators separated by 400 bp; i.e. ergodicity is attained with recordings of no less than 60 min. This is due to the inherent stochastic nature of protein-mediated looping. If, however, DNA molecules are supercoiled, the reduced dimensionality modulating juxtaposition of the protein binding sites accelerates dynamics such that all DNA tethers exhibit similar looping probability and the statistical behavior of the ensemble can be revealed in much shorter observations of a single molecule. Indeed, analysis of looping probability in records for unwound DNA show that a cumulative distribution of looping probabilities equivalent to that attained in 20 min-long measurements is achieved in just 12 minutes (Figure 5B, Supplemental Figure S10). Thus, looping dynamics in supercoiled DNA are deterministic within the time scale of the doubling time of the bacteria, effectively the cell cycle.

**Figure 5.**
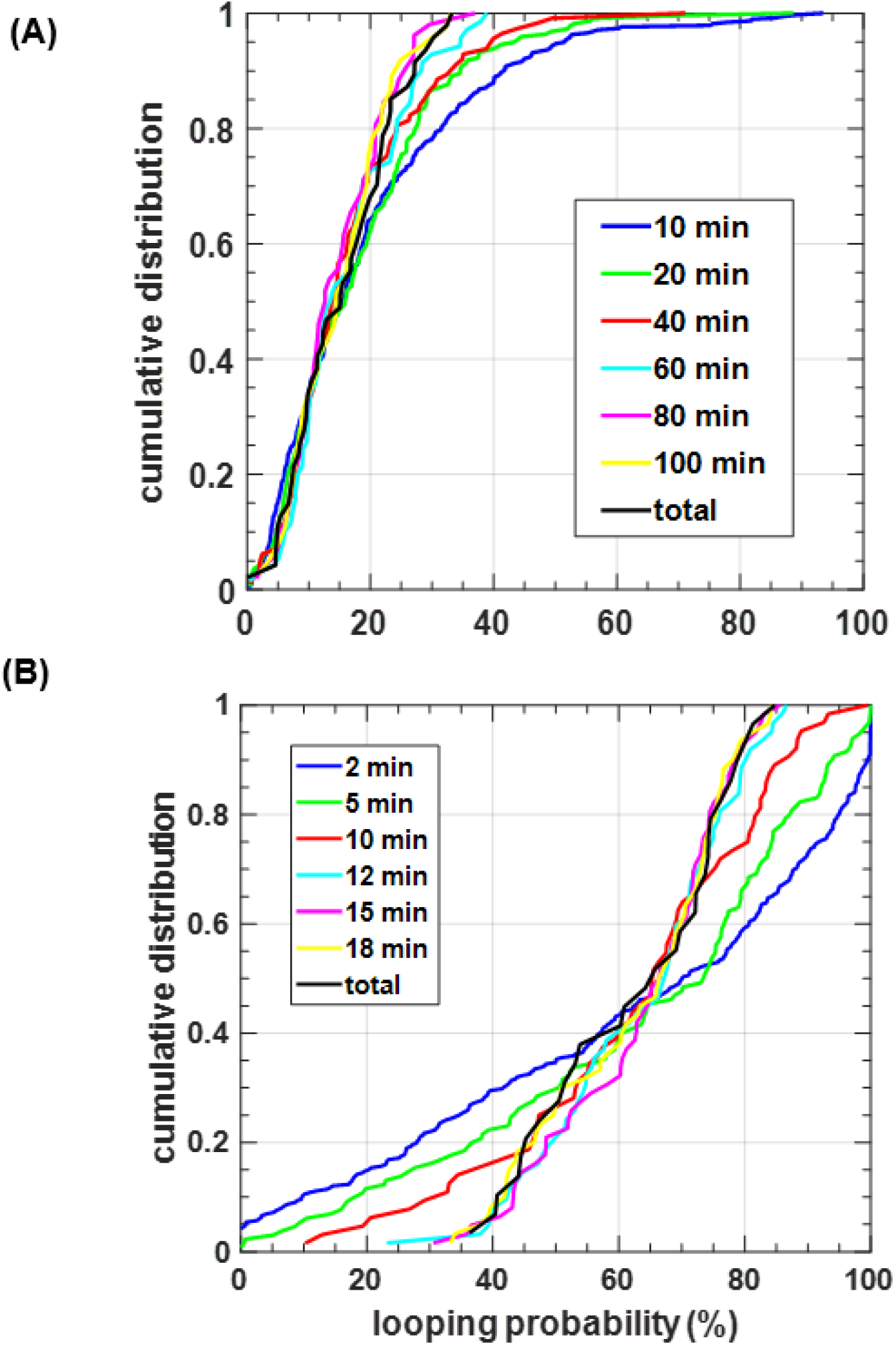
Minimum observation times for ergodicity. Cumulative distributions of the looping probability for (**A**) different periods of TPM observations (10, 30, 40, 60, 80, 100 min) and (**B**) different periods of MT observations (2, 5, 10, 12, 15, 18 min) are plotted along with the cumulative distribution for the entire set of observations (black curves). All records used in this analysis were for 0.25 pN tension and 1.5% supercoiling. Only when observations last at least 60 min (*A*) or 12 min (*B*) do the cumulative distributions of the looping probabilities for individual molecules resemble the distributions for the maximum length observations. With shorter observation periods, the looping probabilities measured for individual molecules varies widely. For example, in the TPM data in *A*, looping probabilities measured for molecules observed for 10-minute intervals (blue) displayed a significant fraction of high probabilities that did not appear for the molecules observed for 60 min or more, up to 5 hours, (black)). In *B*, looping probabilities spanning the entire range result for 2- or 5-min periods of observation but narrow to the 40-80% range when observation times are 12 min or longer.

## Conclusion

Recently, LacI was observed to hop along the double helix (54). This feature together with negative supercoiling is probably key for the protein to efficiently locate a binding site, contact a secondary binding site, and maintain a constant ratio of looped versus unlooped states in the various states of tension and torsion that likely develop during the cell cycle. This ratio gives a definitive regulatory outcome to looping based on inherently stochastic molecular activity. Additional parameters may contribute to achieving a specific outcome, but for straightforward protein-mediated looping, DNA supercoiling appears to be key.

## Acknowledgments

We are grateful to Todd Lillian at the University of Southern Alabama for critical discussions. This work was supported by grants from the National Institutes of Health to L.F. (R01 GM084070) and F.L (1R21AI125973-01A1). Kathleen Matthews at Rice University generously provided the LacI protein. Plasmid pRLM118 was a gift from Roger McMacken (Johns Hopkins University).

## Author Contributions

Conceptualization, D.D., L.F., Y.Y., S.K.; Methodology, S.K., Y.Y., W.X., D.D., F.L.; Software, S.K.; Formal Analysis, Y.Y., W.X., S.K., A.Z.; Investigation, S.K., Y.Y., A.Z., W.X.; Writing – Original Draft, L.F., Y.Y., W.X., A.Z.; Writing, Reviewing, Editing, L.F., D.D.; Funding Acquisition, L.F., F.L.; Resources, F.L.

## Data availability statement

Data for the figures presented in this article are deposited in Emory Dataverse, https://doi.org/10.15139/S3/YTKRI8

## Material and Methods

### Preparation of Proteins and DNA constructs

LacI was provided by Kathleen Matthews (Rice University). *E. coli* HU protein was overexpressed in *E. coli* strain BE257recA (*C600 leu*, *pro*, *lac*, *tonA*, *str*, *recA*) carrying plasmid pRLM118 where the hupA and hupB genes are under control of the lambda PL promoter. To express and purify *E. coli* HU protein, the *E. coli* strain BE257recA/pRLM118 was grown in 4 liters of terrific broth containing 50 μg/mL of ampicillin at 30 °C with vigorous aeration to OD595 reaching ~0.7, and then switched to 42 °C by adding 500 mL of 66 °C terrific broth per liter and grown for an additional 50 min at 42 °C. The cells were harvested by centrifugation at 6,000 rpm 4 °C for 15 min. The supernatant was discarded, and the cell pellet was resuspended in a cold cell lysis buffer (25 mM HEPES-KOH, pH 7.6, 1 M KCl, 1 mM DTT, 0.25 mg/mL lysozyme, and 1 mM PMSF). The cells in the cold buffer were incubated on ice for 1 hour, then frozen in liquid nitrogen, and stored in a −80 °C freezer. On the following day, the *E. coli* cells were thawed at a 4 °C water bath. After the thawing, the cell lysate was sonicated on ice six times at 30 W with 2 min interval between each sonication and centrifugated at 18,000 rpm for 60 min. The supernatant was dialyzed against buffer A (25 mM HEPES-KOH, pH 7.6, 50 mM NaCl, 1 mM DTT, 0.25 mg/mL lysozyme, 10% glycerol, and 1 mM PMSF) overnight and then loaded onto a 40 mL SP Sepharose FF column equilibrated with buffer A. The HU protein was eluted with a 300 mL NaCl gradient of 0.1-0.6 M NaCl in buffer A. The peak fractions were pooled, dialyzed against buffer A overnight, and loaded onto a DNA-cellulose column. The HU protein was eluted with a 300 mL NaCl gradient of 0.1-0.6 M NaCl in buffer A. The purity of *E. coli* HU protein was monitored using 20% SDS-PAGE and the concentration was determined using the Lorry assay (Biorad). The purified *E. coli* HU protein is free of nuclease contaminations as determined in the Leng laboratory.

DNA constructs (Supplemental Figure 1) were built similarly to others previously described (36). All DNA fragments for TPM experiments were amplified by PCR using pO1O2 (55) or pZV_21_400 (56) plasmids, as a templates, digoxigenin- and biotin-labeled sense and anti-sense primers (Integrated DNA Technologies, Coralville, IA, or Invitrogen, Life Technologies, Grand Island, NY, USA) (Table S1), dNTPs (Fermentas-Thermo Fisher Scientific Inc., Pittsburgh, PA, USA) and Taq DNA Polymerase (New England BioLabs, Ipswich, MA, USA). The final 831bp-long DNA amplicons contained centrally located O1 and O2 operators separated by 400 bp (O1-400-O2 DNA; Supplemental Figure S1A).

DNA tethers used in the MTs measurements were built by amplifying a 3352, 2115 or 2011 bp-long main fragment containing the O1-400-O2 segment in the middle (Supplemental Figure S1B) and using T7 ligase (New England Bio-Labs, Ipswich, MA, USA) to attach ~150 bp-long multiply digoxigenin-, or biotin-, labeled DNA fragments at opposite ends. These termini were necessary to securely attach one end of the main DNA fragment to an anti-digoxigenin-coated, flow-chamber surface and the other end to a streptavidin-coated bead.

The main fragments were amplified with a dNTP mix (Fermentas-Thermo Fisher Scientific Inc., Pittsburgh, PA, USA) and a forward primer containing an ApaI restriction site paired with a reverse primer containing an XmaI restriction site. A double digestion of the amplicon with ApaI and XmaI was purified prior to ligation. ~150 bp biotin- or digoxigenin-labeled DNA anchor fragments were generated by ApaI or XmaI restriction in the middle of 302 bp PCR amplicons produced using dATP, dCTP, dGTP, dTTP (Fermentas-Thermo Fisher Scientific Inc., Pittsburgh, PA, USA), and digoxigenin-11-dUTP (Roche Life Science, Indianapolis, IN, USA) or biotin-11-dUTP (Invitrogen, Life Technologies, Grand Island, NY, USA) in a molar ratio of 1:1:1:0.7:0.3. The product of ligation of the main fragment and two anchor fragments was purified and the total variation in DNA tether length, due to random placement of 30% biotin- or digoxigenin along the anchor fragments did not exceed 50 bp (16 nm) (36) (Supplemental Figure S2).

Details about template and primers that were used to generate the TPM and MT DNA constructs are summarized in Supplemental Table 1.

### Microchamber preparation

Parafilm gaskets were fashioned with a laser cutter (Universal Laser Systems, VLS 860, Middletown, CT) supported on microscope slides, and coverslips were placed on top. Each gasket included inlet and outlet reservoirs positioned just beyond the edges of the coverslip and connected through narrow inlet and outlet channels to the central observation area (Supplemental Figure S3). The assembly was heated briefly on a hot plate at the minimum setting to seal the components together. The narrow channels reduced evaporation of buffer while non-homogeneous flow through the triangular shape of the observation area produced a gradient of tether densities. DNA tethers in *λ* buffer (10 mM Tris-HCl (pH 7.4), 200 mM KCl, 5% DMSO, 0.1 mM EDTA, 0.1 mg/mL α-casein (Sigma-Aldrich, St. Louis, MO) were introduced into the chamber and attached through a single digoxigenin (TPM), or a multiply digoxigenin-labeled tail (MT) to the coverslip coated with anti-digoxigenin (Roche Life Science, Indianapolis, IN, USA). The opposite end of the DNA was attached to a streptavidin-coated bead via a single biotin (TPM), or a multiply biotin-labeled tail (MTs).

The beads used in TPM measurements were 0.32 μm diameter, streptavidin-coated polystyrene beads (Spherotech, Lake forest, IL, USA), while the beads used in MT measurements were 1.0 μm diameter, streptavidin-coated super-paramagnetic beads (Dynabead MyOne Streptavidin T1, Invitrogen, Grand Island, NY)

### Tethered particle motion (TPM) experiments

All TPM experiments were conducted in *λ* buffer at room temperature. The LacI protein was introduced into the chamber before recording ~30 min or longer. For long experiments (up to 5 hours), the inlet and outlet of the chamber were sealed with grease after adding the LacI protein to avoid evaporation.

A Leica DM LB-100 microscope (Leica Microsystems, Wetzlar, Germany) with an oil-immersion objective (63X, NA 0.6–1.4) and differential interference contrast optics was used to observe tethered beads with a CV-A60 video camera (JAI, Copenhagen, Denmark). The absolute XY positions of each bead was recorded at 50 Hz with a custom Labview (National Instruments, Austin, TX, USA) program. Vibrational or mechanical drift in the position of each tethered bead was removed by subtracting the average position of multiple, stuck reference bead(s) within the same field of view (38, 57). The excursion of each tether was then calculated as 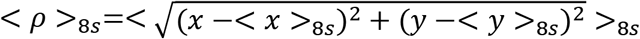, in which < *x* >_8s_, and < *y* >_8s_ are eight-second moving averages representing the coordinates of the anchor point of a bead. Changes in the excursion of the bead reflect conformational (length) changes of the DNA tether (39, 58, 59).

The beads that exhibited (*x*, *y*) position distributions with a ratio of the major to minor axes greater than 1.07, were discarded, since they were likely to be tethered by multiple DNA molecules (38). The excursion data from the time records of the beads, in the same experimental conditions, which passed this “symmetry test” were pooled to generate probability distribution histograms. These summarized the average excursion distribution and, in the presence of LacI, they indicated three excursion peaks (two looped states and one unlooped state) (36). The histogram of each selected temporal trace was fitted with three Gaussians, then the looping probability was calculated by dividing the area under the Gaussians corresponding to the two looped states (peaks with shorter excursion) by the total area under all three Gaussians. The mean value of looping probability under each protein condition were weighted by the length of each trace.

### Magnetic tweezer experiments

The permanent magnets used in magnetic tweezer can move vertically along and rotate around the optical axis of the microscope to change the tension, or torsion, on the beads, and thus, twist the DNA tethers (Supplemental Figure 4 left panel). The hardware, as well as the bead tracking algorithms, have been previously described(56). The *x*, *y*, *z* and *t* coordinates of tethered beads and at least two stuck beads were recorded after calibration. Stretching DNA tethers under high tension (~2 pN) extends a 2115 or 2011 bp DNA template to ~0.7 or 0.67 *μm*, which was used to identify tethers with acceptable contour lengths. Twisting DNA tethers to measure the “hat curve” (Supplemental Figure S4 right panel) helps to identify and discard beads tethered by multiple DNA molecules. Selected tethers were recorded for 3 minutes under three different levels of tension (0.25, 0.45 or 0.75 pN) and at a series of twist settings to determine the DNA extension in each condition before adding LacI protein. After introducing 1 nM LacI protein, *x*, *y*, *z* and *t* data were recorded for 20 min at each selected tension and twist setting. The DNA extension *vs*. time data were then analyzed to identify probable looping events and calculate looping probability.

The position versus time records for each bead were taken holding the magnet at an integral number of turns. However, since there were multiple number of tethers in a field of view, the torsionaly relaxed state of each DNA tether was not synchronized with the initial “zero” rotation of the magnet. Thus, the torsionaly relaxed state of DNA molecule was obtained by fitting the extension versus twist curve for each molecule with a parabola (Supplemental Figure S5). Then the effective number of turns applied to a DNA molecule was calculated by subtracting the center of hat curve from the turns introduced with the magnet. Then the supercoiling density was calculated as the effective number of turns divided by the twist of a torsionaly relaxed DNA (number of base pairs/helical pitch).

### Looping probability distributions as a function of observation periods

For individual tethers, the duration of looped (*τ_l_*) and unlooped (*τ_u_*) states was determined using the total variation denoising (TVD) algorithm (60), and the looping probability was calculated as *P* = ∑*τ_l_* /(∑*τ_l_* + ∑*τ_u_*). When the time series was divided into shorter intervals, the looping probability was calculated as: *P_l,T_* = ∑_*T*_*τ_l_*/(∑_*T*_*τ_l_* + ∑*_T_τ_u_*), where *∑_T_* refers to the sum within a time interval, *T*. Five-hour-long TPM measurements were sub-divided into intervals of 10, 20, 40, 60, 80, or 100 minutes. Twenty-minute-long magnetic tweezer records were sub-divided into intervals of 2, 5, 10, 12, 15, or 18 minutes. Distributions of *P_l,T_* and *P* were processed with the “ecdf” function of Matlab to obtain the empirical cumulative distribution of looping probabilities for each time interval.

