## Supplemental Figures for "Negative DNA supercoiling makes protein-mediated looping deterministic and ergodic within the bacterial doubling time" (from another version of this preprint: version 1, 2021-02-26)

Figure S1

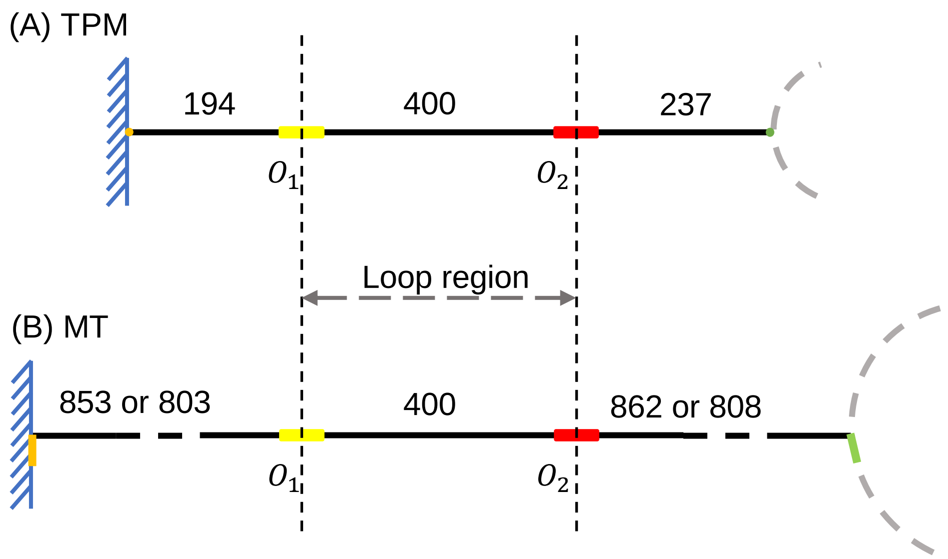


**Figure S1. Schematic diagrams of the DNA constructs used in TPM and MT measurements.** Both constructs have the same central 400 bp loop region segment flanked by the O1 operator, toward the attachment to the glass surface of the microchamber (left, hatched) and O2 operator, toward the bead (right dashed arc). Lengths are given in base pairs.

Figure S2

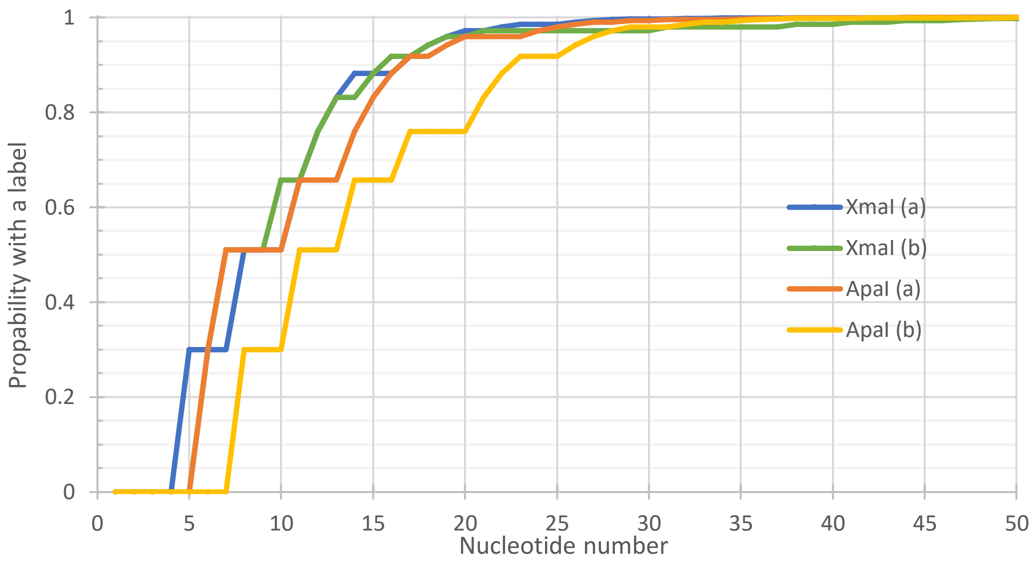


**Figure S2. The probability of labeling along the anchorage fragment as a function of the distance from the ligation junction with the main fragment.** The anchorage fragments are prepared by digesting with XmaI or ApaI approximately 300 bp-long, biotin- or digoxigenin-labeled DNA amplicons centered on the multi-cloning site of pBluescriptKS+. Since the labels are attached to dUTP and the sequences are known, the probability of incorporating labeled nucleotides can be calculated as a function of the distance from the ligation junction (x) as the complement of the probability of not incorporating any label within that span. Since the fraction of labeled nucleotides was 0.3, the probability of labeling at each adenine nucleotide was 0.3. The probability that not even one label would have been incorporated within a segment x nucleotides long was $P_{unlabeled} = 0.7^{N_{A,x}}$ , where $N_{A,x}$ is the number of adenines in the segment. Then the probability that at least one label will be incorporated in the segment is $P_{labeled} = 1 - 0.7^{N_{A,x}}$ . There is about 0.95 probability that at least one label will be incorporated within 27 bp from the junction. The variation of the effective tether length due to labeling will range from 14 to 49 bp or about 12 nm.

Figure S3

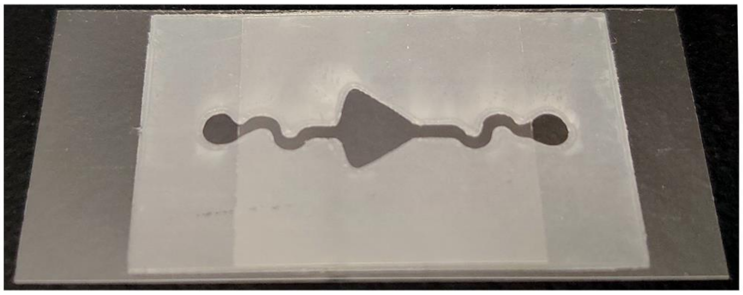

**Figure S3. An image of a triangular microchamber**. A parafilm gasket cut with a laser cutter was placed between cleaned 24 X 50 mm and 22 X 22 mm coverslips and heated on a benchtop heater on a low setting for a few seconds to seal the assembly.

Figure S4

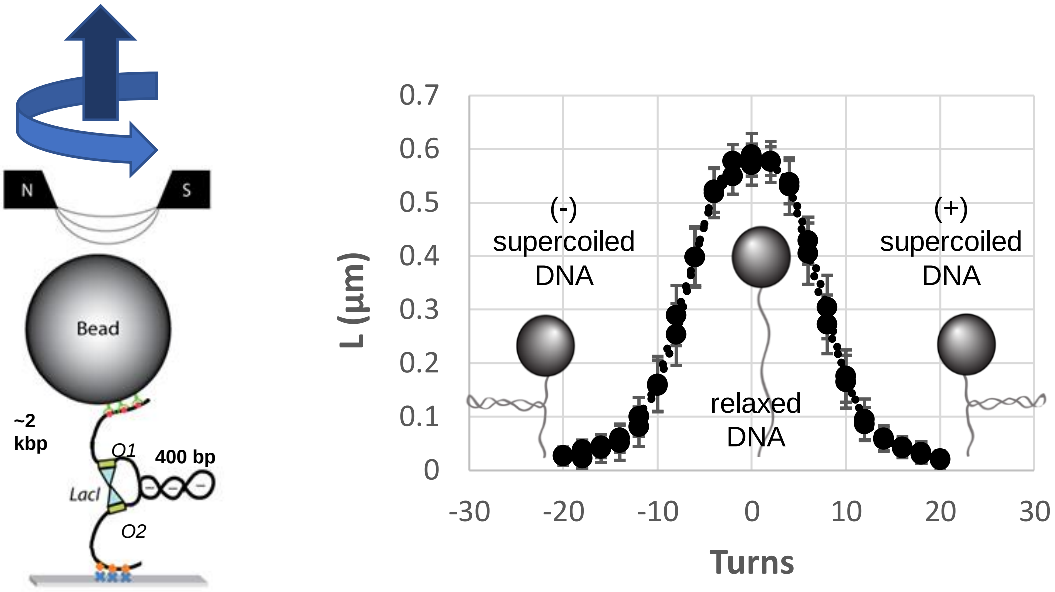


**Figure S4. DNA supercoiling manipulation and looping by magnetic tweezers.** DNA supercoiling refers to the over- or under-winding of a DNA molecule, which is often quantified as the ratio of the number of turns added, or subtracted, to a DNA molecule with respect to the number of helical turns in the torsionally relaxed state.

(Left) Magnetic tweezers are a convenient tool with which to modulate DNA supercoiling *in vitro*. A magnetic field within a microchamber attracts and orients a super para-magnetic bead attached to a single DNA molecule, applying tension to the DNA (dark, vertical arrow) and, upon rotation, twisting the DNA (light, circular arrow). For supercoiling, a DNA molecule must be anchored to the surface and the bead through multiple attachments.

(Right) Twisting a 2115 bp DNA molecule under 0.3 pN of tension induces plectonemes that decrease the overall extension of the torsionally relaxed DNA (0 turns) as shown in this typical extension versus twist plot.

### Figure S5

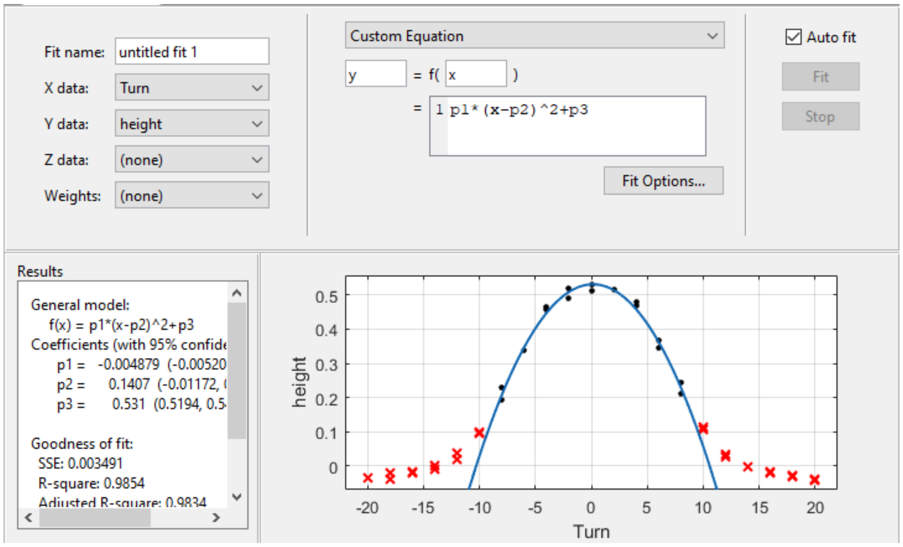


| | | Turn | CorrectTurn | Supercoil D | Ploop |
|---|---|---|---|---|---|
| | | -1 | -1.1407 | -0.5609116 | 0 |
| Center | 0.1407 | -2 | -2.1407 | -1.0526374 | 0 |
| Height | 0.531 | -3 | -3.1407 | -1.5443631 | 0 |
| | | -4 | -4.1407 | -2.0360889 | 4.918486 |
| | | -5 | -5.1407 | -2.5278147 | 42.32796 |
| | | -6 | -6.1407 | -3.0195404 | 93.48375 |
| | | -7 | -7.1407 | -3.5112662 | 98.72791 |

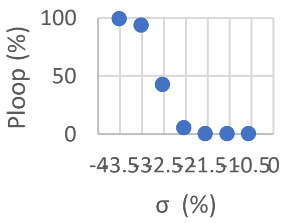


**Figure S5. Finding the center of a twist versus extension curve.** The number of turns added to a DNA molecule was calculated by fitting the extension versus twist curve with a parabola to determine the center.

Figure S6

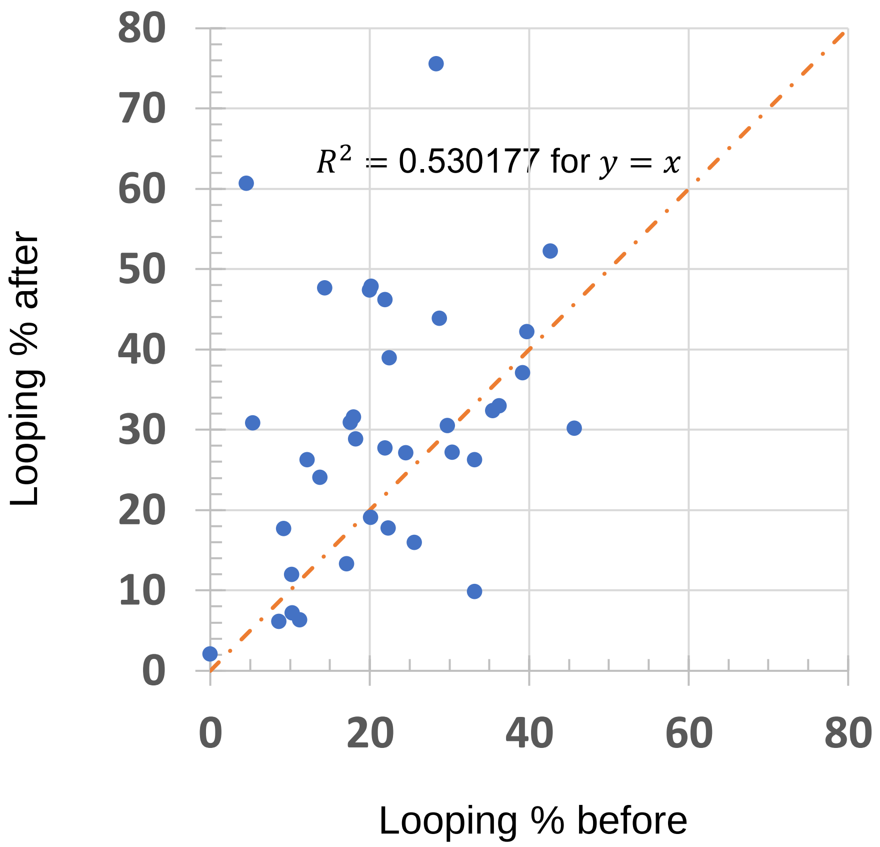


**Figure S6. Looping % before and after LacI replacement**
Using tethered particle motion analysis, the percentage of time spent in the looped state was measured for a field of view of individual tethers in a first observation period. Then LacI was washed out and re-introduced. Following that the percentage of time spent in the looped state for the same set of tethers was assessed for a second observation period. Fitting the data with $y = x$ yields $R^2 = 0.530177$. This value of $R^2$ indicates that there was no correlation between the looping percentages measured for individual tethers in the two observation periods indicating that the percentage of looping is not conditioned by a given DNA molecule.

### Figure S7

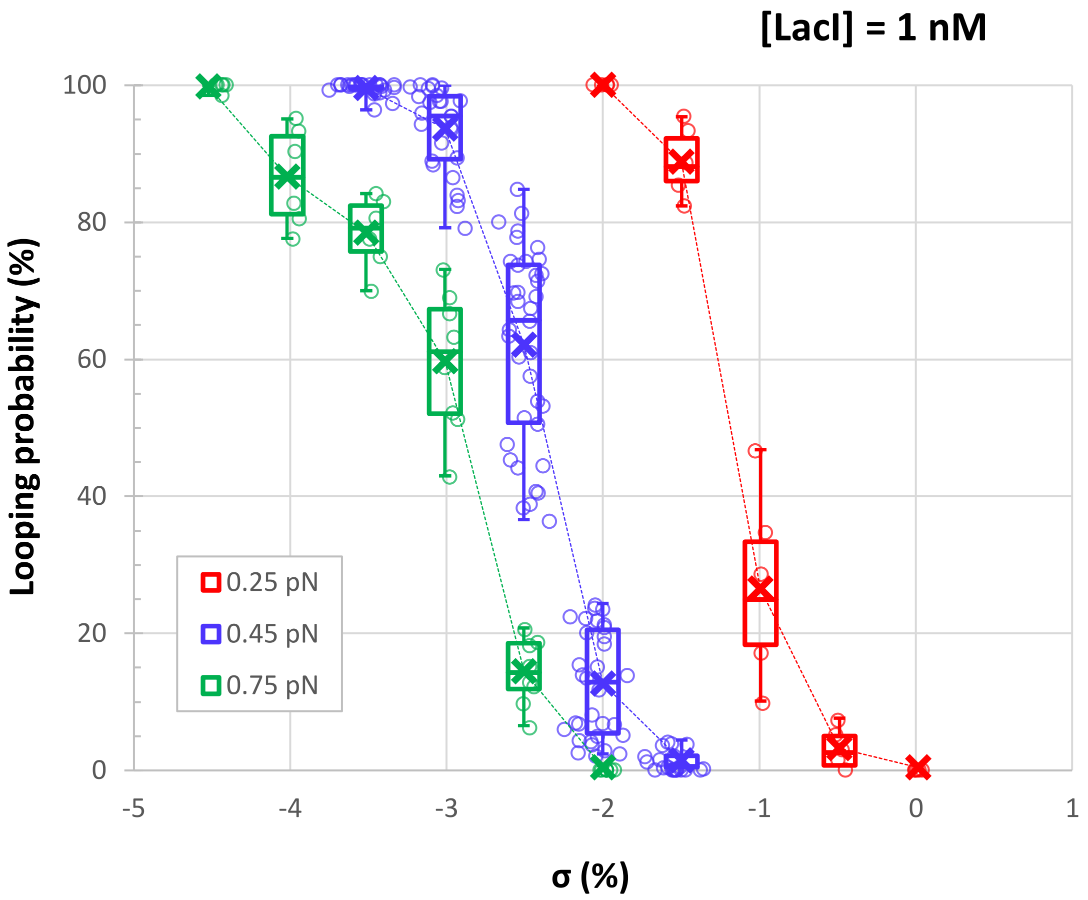


**Figure S7. Low variation is observed for looping percentages measured in supercoiled DNA.** A box-whisker plot summarizes the looping probabilities of DNA tethers in the presence of 1 nM LacI, under 0.25, 0.45 or 0.75 pN of tension and a range of negative supercoiling values (σ). The midline of the box indicates the median and the cross the mean of each distribution. The upper and lower borders of each box indicate the upper and lower quartiles of the measured looping probabilities. The whiskers reveal the complete range of the data. Supercoiling greatly restricted the ranges of data compared to corresponding data for DNA tethers that were not subject to mild tension and torsion as shown in Figures 1 and 2 of the main article.

Figure S8

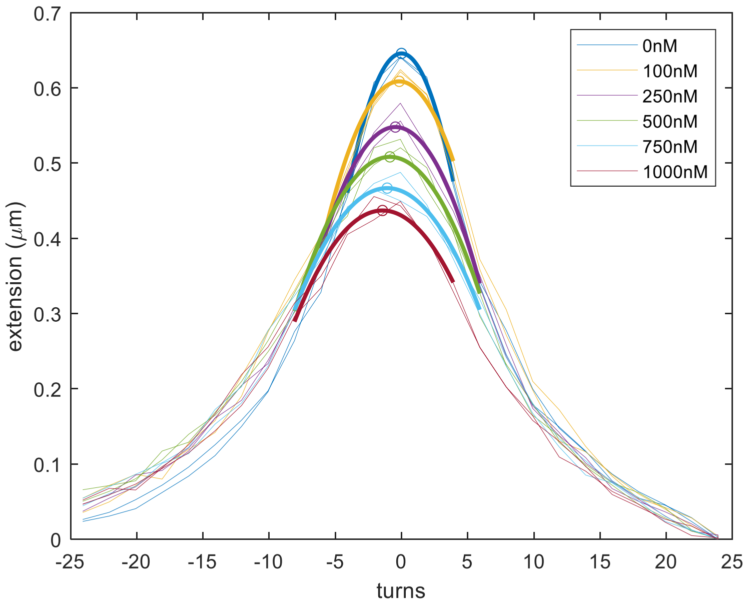


**Figure S8. HU binding significantly contracts but mildly unwinds DNA.** A single 3352 bp DNA tether under 0.45 pN of tension in a magnetic tweezer was repeatedly wound and unwound in a buffer containing 200 mM KCl with various concentrations of HU ranging from 0 to 1000 nM. Increasing HU concentrations steadily reduced the extension of the DNA tether by nearly 30% but produced only very mild negative supercoiling. A parabolic curve (thick) was fit to each pair of winding and unwinding curves corresponding to different concentrations of HU. The vertex of each parabola (open circle) represents the shift of the twist versus extension curve due to the supercoiling induced by HU protein binding to the DNA tether.

### Figure S9

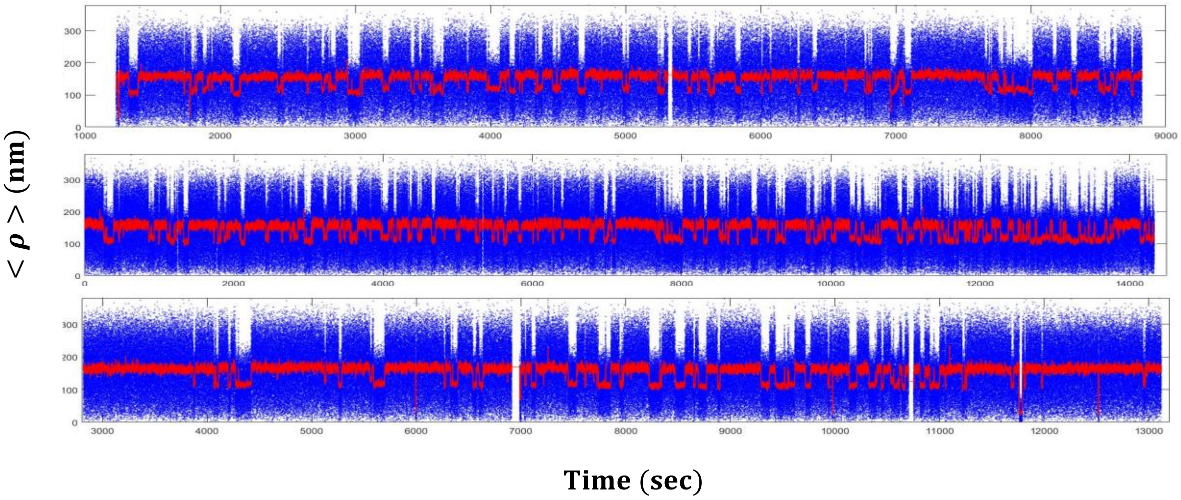


**Figure S9. Long TPM recordings.** Three representative recordings are shown from a set of fifty lasting over one hour. The blue dots represent the two-dimensionally projected amplitude of momentary excursions of the tethered bead; the red trace is the moving, 8-second time average of these values.

### Figure S10

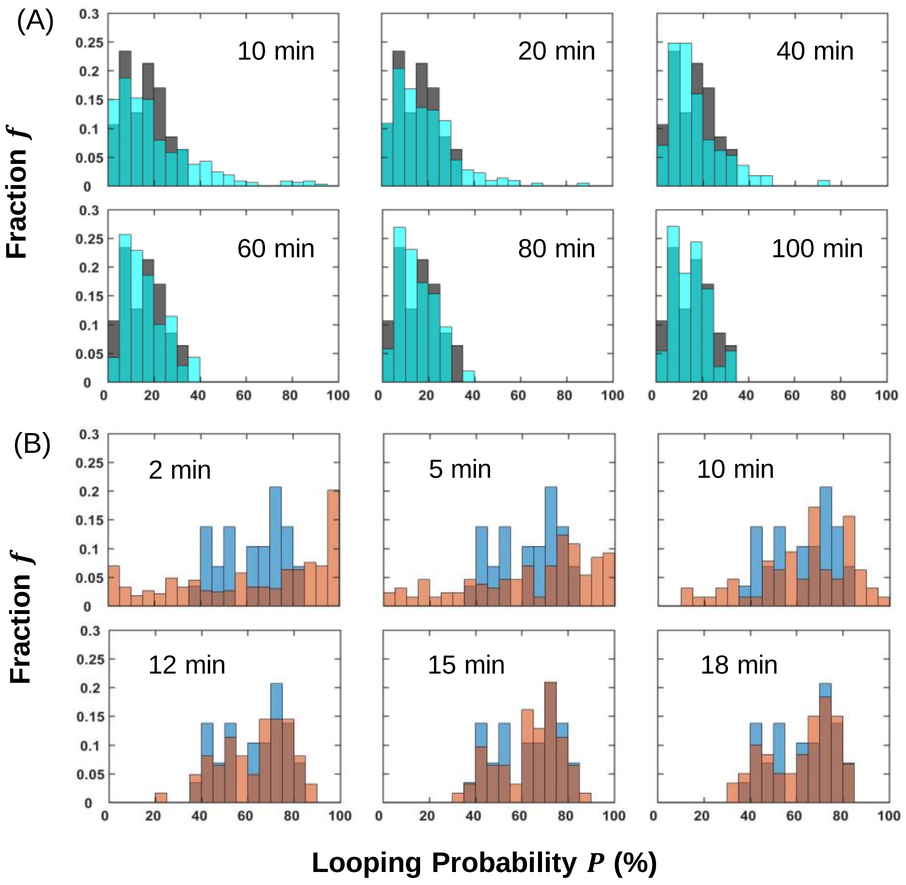


**Figure S10. Sufficiently long observations are ergodic.**
(A) Distributions of the looping probabilities derived from TPM records lasting 5 hours (black) or split into shorter segments as indicated (cyan). (B) Distributions of the looping probabilities derived from MT records lasting 20 min (blue) or split into shorter segments as indicated (brown). All data was recorded at 0.45 pN of tension and -1.5% supercoiling. The distributions of looping probabilities for the corresponding full-length record is included for comparison in each panel.

### Table S1

| Construct | Template | Sense primer | Anti-sense primer | Restriction enzymes |
|---|---|---|---|---|
| O1-400-O2 (TPM) | pO1O2[1] | D-S/pO1O2_401/1588 | B-A/pO1O2_401/2418 | none |
| O1-400-O2 (MT) | pO1O2, pZV_21_400[2] | S/pO1O2_401/929_XmaI, S/JBOIDO1_400/1997_ApaI | A/pO1O2_401/3043_ApaI, A/JBOIDO1_400/3810_XmaI | XmaI, ApaI |
| 3352 bp segment (MT) | pYY_I1_400_BstEII [3] | S/JBOIDO1_400/1783-Apa1 | A/JBOIDO1_400/5113-Xma1 | XmaI, ApaI |
| Bio-, dig-tails | pBluKSP+ | A/pIT_Loop3/1665 | Pol1979R | XmaI, ApaI |

### Table S2

| Primer name | Sequence |
|---|---|
| D-S/pO1O2_401/1588 | [dig]-tgctcgcttcgctacttg |
| B-A/pO1O2_401/2418 | [bio]-tgactgggttgaaggctc |
| S/pO1O2_401/929_XmaI | tgCCCGGGacccggaaagacatgc |
| A/pO1O2_401/3043_ApaI | ctGggCCCggtgaatccgttagcga |
| S/JBOIDO1_400/1997_ApaI | acccgtGGGCCCagcatcctctctcgtttcatc |
| A/JBOIDO1_400/3810_XmaI | gcagcgCCCGGGtgagcgaggaagcggaagag |
| A/pIT_Loop3/1665 | ggcgattaagttgggtaacg |
| Pol1979R | tgtggaattgtgagcggata |
| A/JBOIDO1_400/5113-Xma1 | acaacacccgggATCATGTAACTCGCCTTG |
| S/JBOIDO1_400/1783-Apa1 | atgttcgggcccAGTAACCCGTATCGTGAGC |

**Supplementary Tables of PCR reagents. Table S1** (Top) The DNA constructs were constructed using plasmid templates with the indicated pairs of primers. **Table S2** (Bottom) Sequences for each primer. Bio- or B- is a biotin tag at the 5’ end of the primer, and D- or dig- is a digoxigenin tag at 5’ end of the primer.


[1] G. Fulcrand *et al.*, DNA supercoiling, a critical signal regulating the basal expression of the lac operon in Escherichia coli. *Scientific reports* **6**, 19243 (2016)

[2] Z. Voros, Y. Yan, D. T. Kovari, L. Finzi, D. Dunlap, Proteins mediating DNA loops effectively block transcription. *Protein Science* **26**, 1427-1438 (2017)

[3] Y. Yan, F. Leng, L. Finzi, D. Dunlap, Protein-mediated looping of DNA under tension requires supercoiling. *Nucleic Acids Res* **46**, 2370-2379 (2018)
